# Stem-cell-ubiquitous genes spatiotemporally coordinate division through regulation of stem-cell-specific gene networks

**DOI:** 10.1101/517250

**Authors:** Natalie M Clark, Eli Buckner, Adam P Fisher, Emily C Nelson, Thomas T Nguyen, Abigail R Simmons, Maria A de Luis Balaguer, Tiara Butler-Smith, Parnell J Sheldon, Dominique C Bergmann, Cranos M Williams, Rosangela Sozzani

**Affiliations:** Department of Plant and Microbial Biology, North Carolina State University, Raleigh, NC 27695; Biomathematics Graduate Program, North Carolina State University, Raleigh, NC 27695; Department of Electrical and Computer Engineering, North Carolina State University, Raleigh, NC 27695; Department of Biology, Stanford University, Stanford, CA 94305; Department of Biology, Denison University, Granville, OH 43023; Howard Hughes Medical Institute (HHMI), Stanford University, Stanford, CA 94305

## Abstract

Stem cells are responsible for generating all of the differentiated cells, tissues, and organs in a multicellular organism and, thus, play a crucial role in cell renewal, regeneration, and organization. A number of stem cell type-specific genes have a known role in stem cell maintenance, identity, and/or division. Yet, how genes expressed across different stem cell types, referred here as stem-cell-ubiquitous genes, contribute to stem cell regulation is less understood. Here, we find that, in the Arabidopsis root, a stem-cell-ubiquitous gene, TESMIN-LIKE CXC2 (TCX2), controls stem cell division by regulating stem cell-type specific networks. Development of a mathematical model of TCX2 expression allowed us to show that TCX2 orchestrates the coordinated division of different stem cell types. Our results highlight that genes expressed across different stem cell types ensure cross-communication among cells, allowing them to divide and develop harmonically together.

## Introduction

Stem cells asymmetrically divide to replenish the stem cell and produce a daughter cell that will go on to differentiate into a specialized cell type. Various mechanisms have been proposed for how pluripotency is maintained, such as signaling pathways within the stem cell niche (SCN) that restrict differentiation, predetermined lineages which ensure stem cells are continuously formed, and cell plasticity which allows differentiated cells to revert to a stem-like state^1–4^. However, most of the pathways that have been shown to maintain pluripotency use local mechanisms, such as short-range signaling, DNA methylation, and chromatin remodeling, that only act on the dividing cell and/or the directly adjacent cells^5–8^. There are likely other networks, upstream of these local mechanisms, which are global in nature and allow for cross-communication across different cell populations.

The Arabidopsis root provides an excellent model system for uncovering these global regulatory networks. The root SCN is well-defined and located at the tip of the root, and as the stem cells asymmetrically divide, the differentiated cells are pushed up the root, resulting in a temporal axis where older cells are more shootward and younger cells are more rootward. Crucially, the movement of cells in the root is constrained due to cell walls, and cell-to-cell signals travel via the plasmodesmata, which are small channels in the cell walls^9^. This lack of cell movement coupled with well-defined marker lines that label specific cell populations^10,11^ allows us to study stem cell identity, division, and maintenance in an isolated environment.

Here, we identified genes expressed specifically (in one stem cell type) and ubiquitously (in all stem cell types) that control stem cell division and maintenance in the Arabidopsis root. We first transcriptionally profiled the individual stem cells using spatially well-defined GFP marker lines and found that near half of the stem cell-enriched genes are expressed in only one stem cell type, while the other half are expressed in multiple cell types. We next used Gene Regulatory Network (GRN) inference to predict that there are not only stem-cell-specific gene networks but also an upstream network that regulates all of the different stem cells. Given that most known mechanisms for maintaining stem cell identity and plasticity are local in nature, we focused on identifying genes expressed in all the stem cells (hereinafter referred to as a stem-cell-ubiquitous gene) that regulates aspects of stem cell maintenance. Using our network prediction, we found that TESMIN-LIKE CXC 2 (TCX2), a member of the family of CHC proteins which are homologues of components of the DREAM cell-cycle regulatory complex in animals^12^, is a key regulator of stem cell division. Further, using Ordinary Differential Equation (ODE) modeling, we show that using the dynamics of TCX2 expression we could predict the timing of stem cell division. Our results provide evidence that genes that participate in global regulatory pathways which span many, different cell types are important for controlling stem cell division and maintenance.

## Results

### Stem-cell-type-specific and stem-cell-ubiquitous transcriptional profile control stem cell pluripotency

To understand how and whether stem-cell-ubiquitous genes contribute to cell identity, maintenance, and/or division, we performed gene expression analysis of the stem cells in the Arabidopsis root, as this offers a tractable system given its 3-dimensional radial symmetry and temporal information encoded along its longitudinal axis. To this end, seven root stem cell markers (Figure 1A), as well as a non-stem cell control (i.e., a population of cells from the root meristem excluding most of the stem cells), were used to identify stem cell-enriched genes, and among those, stem-cell-ubiquitous and stem-cell-specific genes, as it has been shown that there is a correlation between expression levels and functionality in specific cell types^1,13^ (Supplementary Figure 1, see Methods). Notably, we found that the expression profiles of our markers together with known stem cell genes, agree with their known expression domains, supporting that our transcriptional profiles are specific to each stem cell population (Supplementary Figure 1). To measure transcriptional differences between the stem cells and the non-stem cells, we next performed a Principal Component Analysis (PCA.) Looking at the top 3 principal components (50.6% of the variation in the data), the PCA shows that the non-stem cell samples (red) are distant from all of the stem cell populations, suggesting that the stem cells have a different transcriptional signature than the non-stem cells (Figure 1B). Accordingly, when we performed differential expression analysis on these data, we found that 9266 (28% of genes) are significantly enriched (q < 0.06 and fold change > 2) in at least one stem cell population compared to the non-stem cells and considered these genes the stem cell-enriched genes (see M&M and Supplemental Table 1). Thus, this approach allowed us to identify core stem cell genes, as functionally important genes are often enriched in the specific cell populations they control^1,13^.

**Figure 1.**
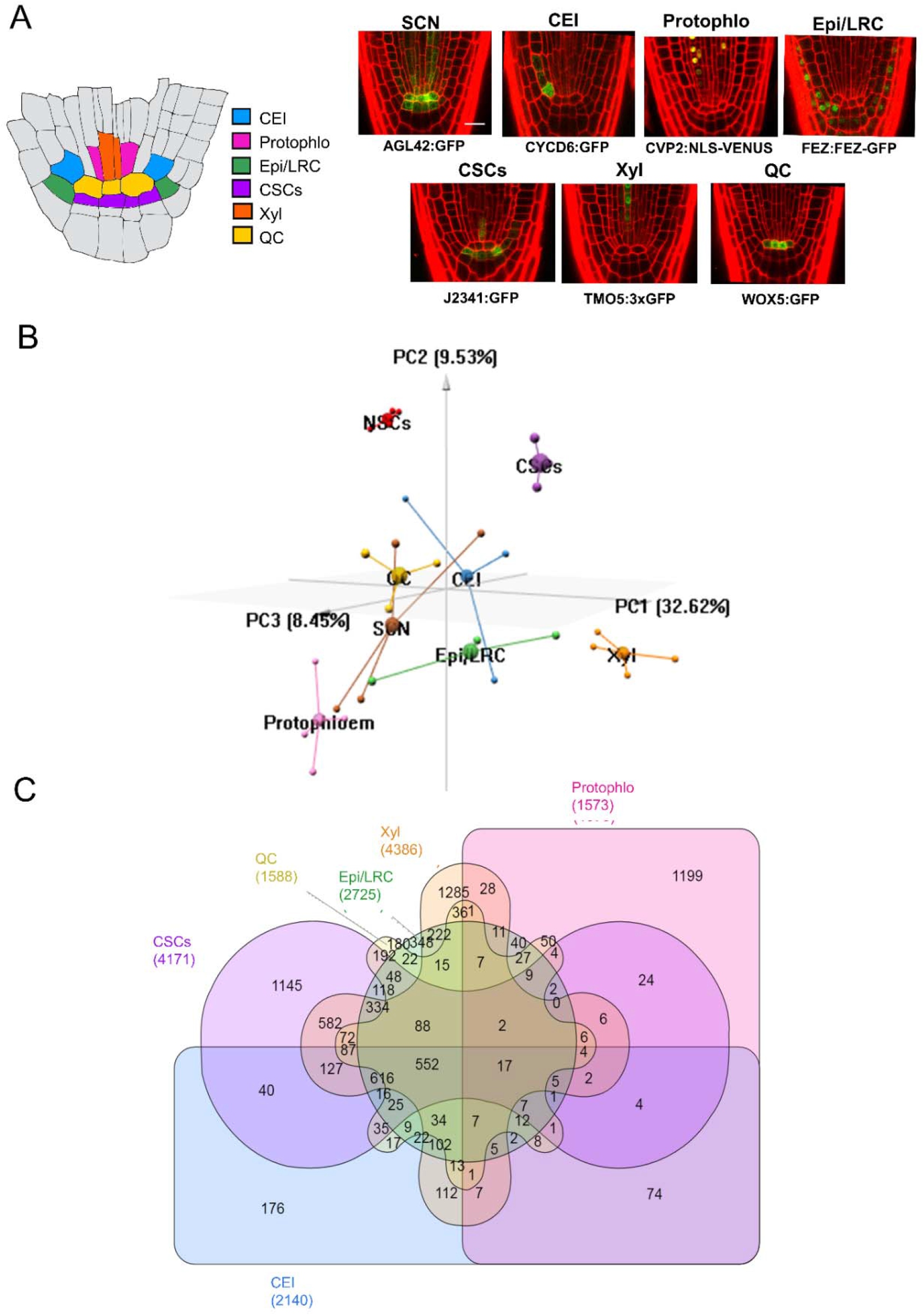
Distribution of cell-specific and cell-ubiquitous genes within the Arabidopsis root stem cell niche. (A) (left) Schematic of the Arabidopsis root stem cell niche. CEI – cortex endodermis initials (blue); Protophlo-protophloem (pink); Epi/LRC – epidermis/lateral root cap initials (green); CSCs – columella stem cells (purple); Xyl – xylem initials (orange); QC – quiescent center (yellow). (left) GFP marker lines used to transcriptionally profile stem cells. SCN – stem cell niche; Scale bar = 20μm. (B) 3D principal component analysis (PCA) of the stem cell transcriptional profiles. The x, y, and z axis represent the three largest sources of variation (i.e. three largest principal components) of the dataset. Small spheres are biological replicates, large spheres are centroids. Red – Non stem cells (NSCs); Brown – SCN; Blue – CEI; Pink – Protophlo; Green – Epi/LRC; Purple – CSCs; Orange – Xyl; Yellow – QC; (C) Distribution of the 9266 stem cell-enriched genes across the stem cell niche. Enrichment criteria are q-value < 0.05 (from PoissonSeq) and fold change in expression > 2.

While the PCA gives us a general idea of how many genes are cell-specific vs cell-ubiquitous, it reduces the dimensionality of the problem to the three largest components of variance. Consequently, we would expect some genes been differentially enriched across all of the stem cell populations. Indeed, when we performed differential expression analysis on the 9266 stem cell-enriched genes (see Methods), we find that 2018 genes (21.8% of the stem cell-enriched genes, hereinafter referred to as the stem-cell-ubiquitous genes) are enriched in at least 4 of the 6 unique stem cell types, with 569 of these 2018 (6.1% of the stem cell-enriched genes) enriched in 5 or 6 cell types (Figure 1C). Moreover, as each stem cell population clusters independently from the others in the PCA, we identified 7248 genes (78.2% of the stem cell-enriched genes), hereinafter referred to as the stem-cell-specific genes, enriched in 3 or less stem cell types, with 4331 of those 7248 genes (46.7% of the stem cell-enriched genes) enriched in only 1 stem cell type. This suggests that each specific stem cell type has its own, unique transcriptional signature.

### Both stem-cell-specific and stem-cell-ubiquitous genes are predicted to be important stem cell regulators

Given the separation between stem-cell-ubiquitous genes and stem-cell-specific genes, we next wanted to know if these two groups of genes have seemingly separated functions or, for example, if stem-cell-ubiquitous genes modulate stem-cell-specific gene expression to orchestrate coordinated processes between different cell types. To test the latter hypothesis, in which stem-cell-specific genes are important for regulating cell type-specific aspects (e.g cell identity), but are regulated by stem-cell-ubiquitous genes so that stem cell maintenance and divisions are tightly coordinated, we used Gene Regulatory Network (GRN) inference and predicted the relationships between all 9266 genes enriched in the stem cells. We developed a machine-learning, regression tree approach to infer dynamic networks from steady state, replicate data (see Methods). Our inferred GRN found regulations among 2982 (32.2%) of the stem cell-enriched genes and predicted that the stem-cell-ubiquitous (red) genes are located in the center of the network, which represents the beginning of the regulatory cascade, and are highly connected to each other (Figure 2A). Meanwhile, the cell-specific (blue) genes mostly regulate each other within the same cell type and are located on the outside of the network, therefore downstream of the cell-ubiquitous genes (Figure 2A). This suggests that the cell-ubiquitous genes are potentially involved in coordinating processes between different stem cells through the regulation of cell-specific genes.

**Figure 2.**
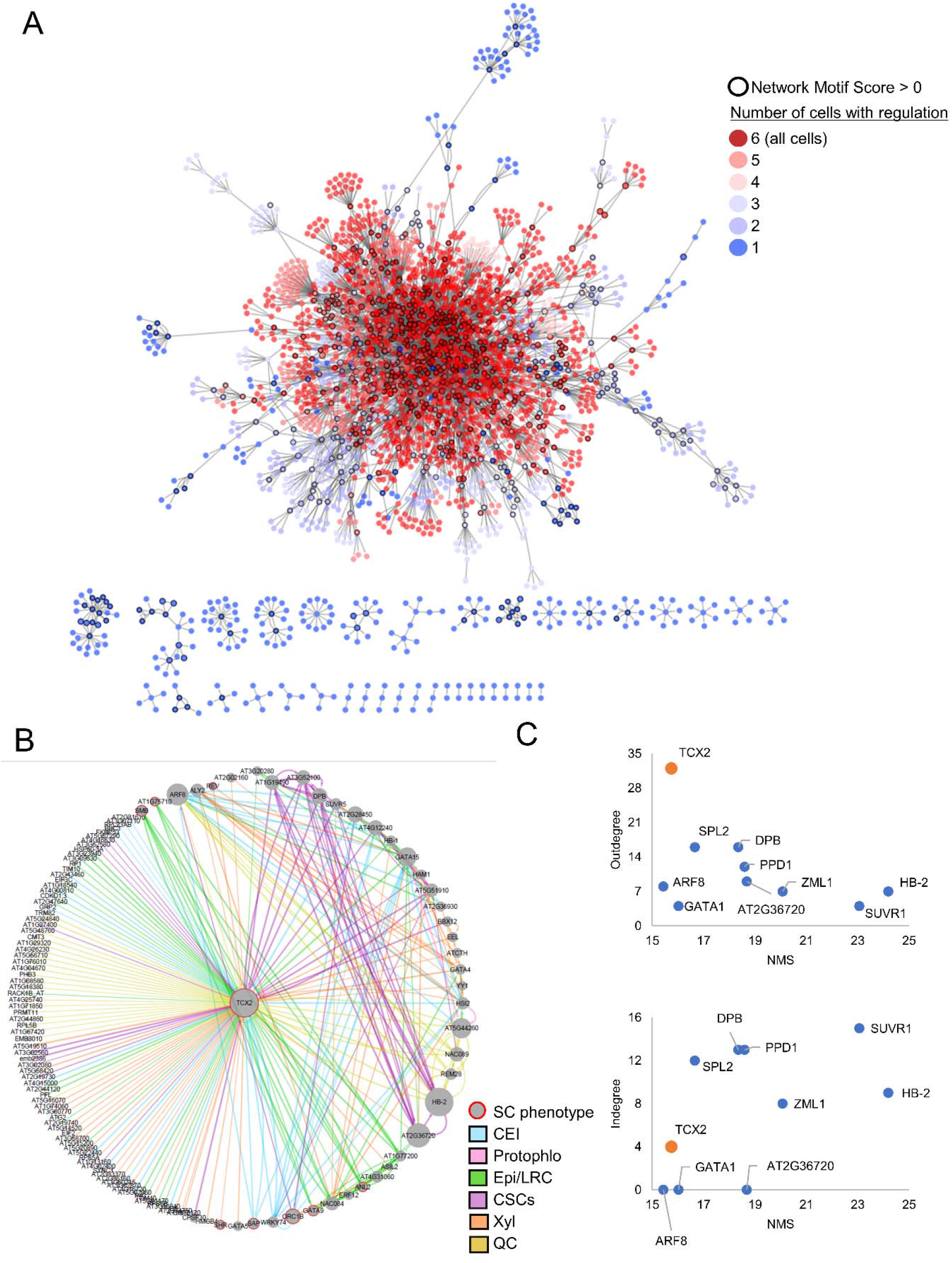
Gene regulatory network (GRN) of the stem cell-enriched genes connects cell-specific and cell-ubiquitous hub genes. (A) Inferred GRN of 2982 out of the 9266 stem cell-enriched genes. Genes are colored based on the number of genes in which they are enriched, with red genes (>3 enriched cells) considered cell-ubiquitous and blue genes (≤ 3 enriched cells) considered cell-specific. Black outlines represent hub genes which have a normalized motif score (NMS) > 0. (B) First-neighbor GRN of TCX2. Gene size represents the NMS score. Red borders represent the genes which have a known stem cell (SC) phenotype. Edge colors represent the cell in which the regulation is inferred. Blue – CEI; Pink – Protophlo; Green – Epi/LRC; Purple – CSCs; Orange – Xyl; Yellow – QC. (C) Outdegree (top plot) and indegree (bottom plot) vs NMS score of the genes with the top 10 NMS scores in (A). TCX2 is highlighted in orange.

We next wanted to identify if the most biologically important genes in the network were cell-specific, cell-ubiquitous, or both, as most results in animals assume that core TFs must be expressed in a cell-specific manner^1^. To predict biological significance, we developed a Network Motif Score (NMS) to quantifies the number of times each gene appears in certain network motifs, such a feedback and feedforward loops (see Methods). These motifs were chosen as they were significantly enriched in our biological network versus a random network of the same size, and have been shown to often contain genes that have important biological functions^14–16^ (Supplementary Figure 2). In our inferred GRN, we found that 737 (24.7%) of the 2982 genes have an NMS > 0, meaning they appear in at least one of the network motifs. To validate the NMS, we found that 22 known stem cell regulators had scores in the top 50% of genes, with 10 of those 22 (45.5%) in the top 25% of genes, supporting that high NMS scores are correlated with stem cell function (Supplemental Table 2). Further, 510 (69.2%) and 217 (31.8%) of these genes are cell-ubiquitous (4 or more enriched stem cells, red) and cell-specific (3 or less enriched stem cells, blue), respectively (Figure 2A). Given that more cell-ubiquitous genes have higher importance scores in our dataset, we focused our downstream analysis on identifying a stem-cell-ubiquitous gene with characteristics of a functionally important regulator.

### TCX2 is an important stem-cell-ubiquitous regulator controlling stem cell division and identity

When we began to examine the stem-cell-ubiquitous regulators, we found that TESMIN-LIKE CXC 2 (TCX2, also known as SOL2), a known homologue of the LIN54 DNA-binding component of the mammalian DREAM complex which regulates the cell cycle and the transition from cell quiescence to proliferation^12,17,18^, had the ninth highest NMS (top 1.2% of genes). This suggests that TCX2 could have an important role across all of the stem cells. To further support the biological significance of TCX2, we examined the subnetwork of its first neighbors (i.e., genes predicted to be either directly upstream or downstream of TCX2). We found that TCX2 is enriched in 5 out of the 6 stem cell types and predicted to regulate at least one gene in all of those cell types, supporting that TCX2 could be a stem-cell-ubiquitous regulator that controls stem-cell-specific core genes (Figure 2B). In addition, when compared to the genes with the top 10 NMS, TCX2 has the highest outdegree (number of edges going out) and low indegree (number of edges coming in), suggesting that TCX2 could orchestrate coordinated stem cell division as suggested by the function of its mammalian homologue^12,17,18^.

If TCX2 is indeed a key regulator for stem cell maintenance and division, we would expect that a change in its expression would cause a developmental phenotype related to these aspects. To test this hypothesis, we obtained two knockdown (*tcx2-1, TCX2-2*) and one knockout (*tcx2-3*) mutants of TCX2, which all show similar phenotypes (Figure 3A, Supplementary Figure 3). Importantly, we observed in *tcx2-3* an overall disorganization of the stem cells, including aberrant divisions in the Quiescent Center (QC), columella, endodermis, pericycle, and xylem cells (Figure 3A). Additionally, *tcx2-3* mutants showed longer roots due to a higher number in meristematic cell/ higher proliferation (Figure 3A, Supplementary Figure 3). Notably, similar phenotypes related to cell divisions have been observed also in the stomatal lineage of *tcx2 sol1* double mutants^12^. To further investigate TCX2’s role in stem cell division, we crossed the cell division (G2/M phase) marker CYCB1;1:CYCB1;1-GFP^19,20^ into the *tcx2* mutant and performed temporal tracking of the GFP signal over time. We first found that average CYCB1;1 expression was higher in the *tcx2* mutant compared to WT. Second, we separated cells expressing CYCB1;1 into 3 categories: low, intermediate, and high expression. We found that significantly more cells in the *tcx2* mutant have high CYCB1;1 expression, while significantly fewer cells have low CYCB1;1 expression. Finally, we calculated the number of consecutive time points each cell shows CYCB1;1 expression. We found that significantly fewer cells in the *tcx2* mutant had 2 consecutive timepoints with CYCB1;1 expression (Supplementary Figure 4). All of these alterations in CYCB1;1 expression in the *tcx2* mutant suggest that reduction of TCX2 expression correlates with more actively dividing cells. Taken together, these results suggest that TCX2, as a stem-cell-ubiquitous gene, regulates stem cell divisions across different stem cell populations.

**Figure 3.**
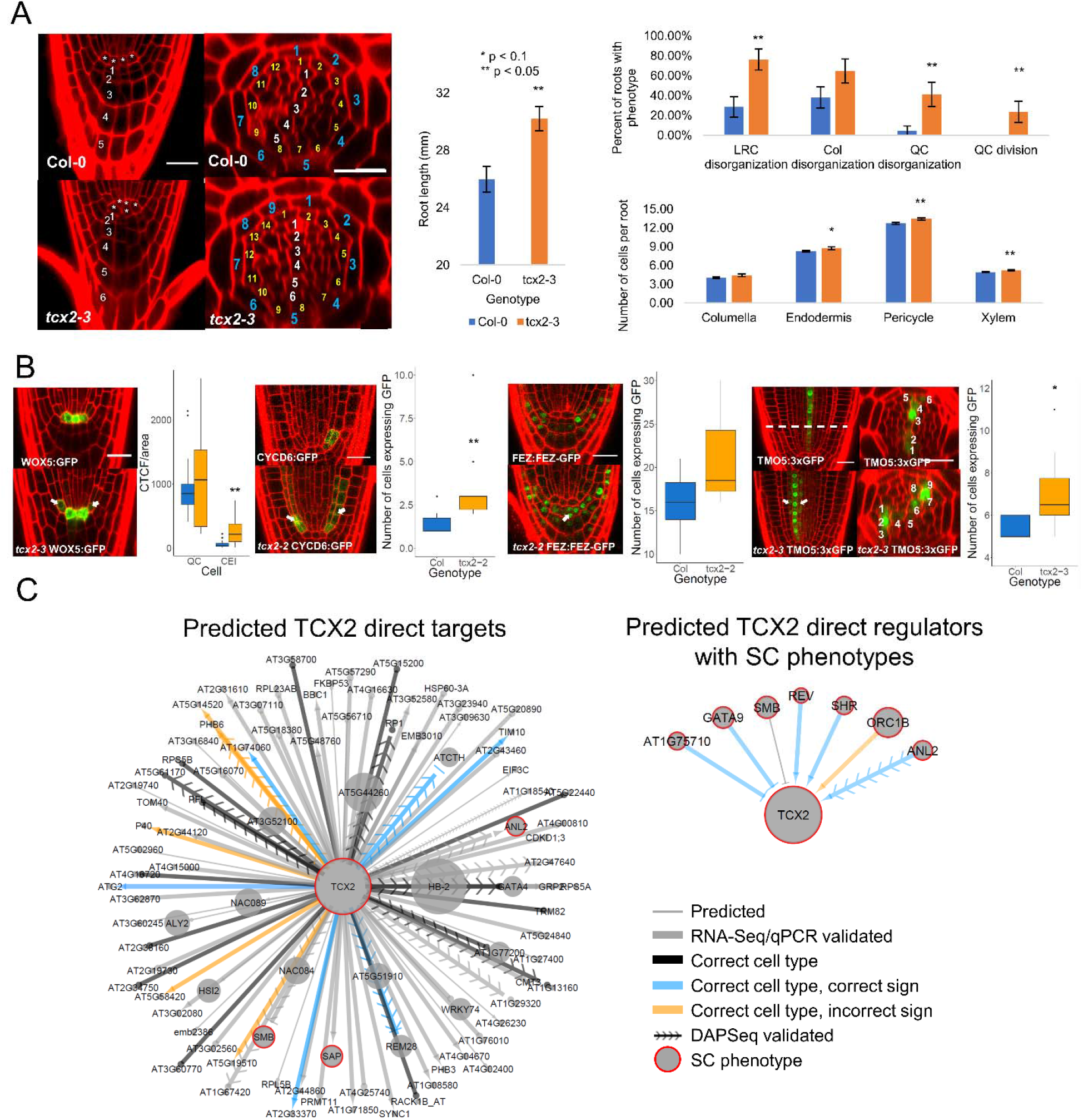
TCX2 controls stem cell division through cell-specific regulators and targets. (A) (Left panel) Medial longitudinal (left) and radial (right) sections of 5 day old WT (top) and *tcx2* mutant (bottom) plants. In medial longitudinal sections, * labels QC cells and numbers denote columella cell files. In radial sections, white numbers denote xylem cells, yellow pericycle, and blue endodermis. (Middle panel) Length of 7 day old WT (blue, n=18) and *tcx2* mutant (orange, n=18) roots. (Right panel) Quantification of stem cell phenotypes (top plot) and number of cell files (bottom plot) in 5 day old WT (blue) and *tcx2* mutant (orange) roots. * denotes p < 0.1, ** denotes p < 0.05, Wilcoxon test. Error bars denote SEM. (B) (left panels) Medial longitudinal sections of 5 day old WOX5:GFP (left), CYCD6:GFP (second from left), FEZ:FEZ-GFP (third from left), and TMO5:3xGFP (right) in WT (top) and *tcx2* mutant (bottom) plants. For TMO5:3xGFP, a radial section (middle) is also shown taken at the location of the white, dashed line. (right panels) Quantification of GFP in WT (blue, n>5) and *tcx2* mutant (orange, n>5) plants. Black dots represent outliers. * denotes p < 0.1, ** denotes p < 0.05, Wilcoxon test. (C) (left) Predicted direct targets of TCX2 and (right) predicted upstream regulators of TCX2 with stem cell (SC) phenotypes. Gene size represents the NMS score. Red borders represent the genes that have a known SC phenotype. Arrows represent predicted activation, bars inferred repression, and circles no inferred sign. Thick edges were validated using qPCR/RNA-Seq. Black edges were predicted in the correct cell type but did not have a predicted sign. Blue edges have the correct cell type and correct sign, while orange edges have the correct cell type but the incorrect sign. Arrows with chevrons are DAPSeq validated. Source data are provided as a Source Data file.

We hypothesized that TCX2 controls stem cell division by regulating important, cell type-specific genes. Notably, all of our stem cell markers, in addition to being expressed only to one stem cell type, are known to have functions in stem cell regulation^21–25^. Thus, we crossed the marker lines for the Quiescent Center (QC; WOX5:GFP), Cortex Endodermis Initials (CEI; CYCD6:GFP), Epidermis/Lateral Root Cap Initials (Epi/LRC;FEZ:FEZ-GFP), and Xylem Initials (Xyl;TMO5:3xGFP) (Figure 1A) into the *tcx2-2* and *tcx2-3* mutant alleles (Figure 3B). Compared to WT, in a *tcx2* mutant the expression pattern of these markers is expanded. Specifically, the QC marker expands into the CEI, the CEI marker expands into the endodermis and cortex layers, the Epi/LRC marker expands into the Columella Stem Cells (CSCs), and the Xyl marker expands into the procambial cells (Figure 3B). This suggests that in the absence of TCX2 coordination of stem cell division and identity is unregulated through an unknown mechanism.

When we examined the predicted upstream regulators and downstream targets of TCX2, we found that 75% are predicted to be cell-specific (expressed in ≤3 stem cell types), suggesting that TCX2 could be regulated and it regulates targets in a cell type-specific manner. (Supplemental Table 3). Thus, to identify additional cell-specific regulators as well as targets of TCX2, we obtained mutants of the transcription factors (TFs) predicted to be TCX2’s first neighbors (i.e. directly upstream or downstream) that also had high NMS scores (Figure 3C, Supplemental Table 3). Two of the genes, SHORTROOT (SHR), and SOMBRERO (SMB) have phenotypes in the stem cells of their loss-of-function mutants, while the loss-of-function mutant of STERILE APETALA (SAP) is homozygous sterile^22,24,26–28^. Additionally, a quadruple mutant of REVOLUTA (REV) together with three other xylem regulators results in missing xylem layers^29^. Further, we obtained loss-of-function mutants of GATA TRANSCRIPTION FACTOR 9 (GATA9), AT1G75710, ORIGIN OF REPLICATION COMPLEX 1B (ORC1B), ANTHOCYANINLESS 2 (ANL2), and REPRODUCTIVE MERISTEM 28 (REM28), which showed root stem cell phenotypes (Figure 3C, Supplementary Figure 5). We were able to validate that TCX2 was differentially expressed (p<0.05) in *gata9, at1g75710, rev, orc1b*, and *anl2* mutants using qPCR as well as in the SHR overexpression line^22^. Further, we performed FACS coupled with RNA-Seq on the 4 marker lines (WOX5:GFP, CYCD6:GFP, FEZ:FEZ-GFP, and TMO5:3xGFP) that we crossed into the *tcx2* mutant to determine the effect of TCX2 on its predicted downstream stem-cell-specific targets. In addition, we performed RNA-Seq on tissue from the stem cell area of the *tcx2* mutant (Supplemental Table 5). Using these data, we were able to validate that 77.78% of the predicted direct targets of TCX2 are differentially expressed in the *tcx2* mutant stem cells. Further, 41.54% of these edges are predicted in the correct cell type, and of those edges predicted in the correct cell type that had a predicted sign, 58.33% of the edge signs are correctly predicted (slightly better than randomly assigning edge signs, which would have a 50% rate of success). To validate some of the direct interactions between TCX2 and its downstream targets, we mined a published DAP-Seq dataset from Arabidopsis leaves^30^ and were able to confirm that TCX2 can directly bind 15.05% of its predicted direct targets (Figure 3C, Supplemental Table 4). Overall, these results suggest TCX2 orchestrates coordinated stem cell divisions through stem-cell-specific regulatory cascades.

### The TCX2 regulatory network changes over time to regulate stem cell division

Given that most of the validated upstream regulators of TCX2 are stem-cell-type specific (Supplemental Table 3), we propose that these cell-specific regulators modulate the dynamics of TCX2 expression in individual cell types. In turn, changes in TCX2 dynamics correlate with changes in expression of its downstream targets (Figures 3C, 3D). Thus, we hypothesized that different dynamics of TCX2 in specific stem cells, as well as changes in TCX2 expression, could be used to predict when each stem cell population divides.

If TCX2 expression is dynamically changing over time in a cell-specific manner, we would predict that the TCX2 GRN also changes temporally. Specifically, we could expect that TCX2 differentially regulates its targets in specific cell types at certain times depending on its expression levels. Thus, to determine if the TCX2 regulatory network changes over time, we first selected 176 genes of interest that were differentially expressed in the *tcx2* root tip sample (Supplementary Table 5) as well as enriched in the stem cells, as these are most likely to be the downstream of TCX2 across different stem cell populations. We inferred GRNs using a time course of the root meristem that is stem cell-enriched (hereinafter referred to as the stem cell time course, see Methods) to predict one network per time point (every 8 hours from 4 days to 6 days). We found that genes in the first neighbor network of TCX2 have different predicted regulations depending on the time point. Specifically, most of the regulation to and from TCX2 are predicted to occur between 4 days (4D) and 5 days (5D), which is the developmental time at which many stem cell divisions take place^22^. (Supplementary Figure 6). Thus, since our gene expression data suggest that loss of TCX2 function correlates with an increase in stem cell division, we hypothesized that most of the TCX2-regulated stem cell division is occurring between 4D 16H and 5D, time at which TCX2 expression decreases at least by 1.5 fold-change (Supplementary Figure 6).

To test how these time- and cell-specific GRNs affect TCX2 expression and therefore cell division, we built a mechanistic model of the GRNs predicted every 8 hours from 4D to 5D (see M&M and Supplementary Information). We used our stem cell time course to determine the cell-specific networks at each time point and constructed equations for each gene in the network (Figure 4A, Supplementary Figure 7). Unlike our GRN, which only predicts the regulations in each cell at each time point, our Ordinary Differential Equation (ODE) model converts the network prediction into a quantitative model of gene expression. Thus, this model allowed us to quantify how TCX2 dynamics change over time and to correlate significant changes in expression with cell division. Our model included the possibility of some of the proteins moving between cell types, as this is a known local signaling/ cell-to-cell communication mechanism^27,31^. Specifically, we used scanning Fluorescence Correlation Spectroscopy (Scanning FCS) and observed that TCX2 does not move between cells, thus suggesting a cell-autonomous function, while observed movement of WOX5^32,33^ and CRF2/TMO3 between cells is in line with a non-cell-autonomous function (Supplementary Figure 8). As our sensitivity analysis predicted that the oligomeric state of TCX2 in the Xyl, diffusion coefficient of WOX5 from the CEI to the QC, and diffusion coefficient of WOX5 from the QC to the Xyl were three of the most important parameters in the model, we experimentally determined these parameters (Supplementary Figure 8, Supplemental Table 6). Given that our network and time course data predict that TCX2-mediated cell division is tightly coordinated and controlled between 4D 16H and 5D, we wanted to ensure that we accurately measured TCX2 dynamics in this time period to produce the best predictive model of stem cell division. To this end, we quantified the expression of the TCX2:TCX2-YFP marker in different stem cells every 2 hours from 4D 18H to 4D 22H (hereinafter referred to as the YFP tracking data) (Figure 4B, see M&M). We then used the average expression of TCX2 in each cell at each time point to estimate parameters in our model (Supplemental Table 7). The result of this model is thus a spatiotemporal map of the expression dynamics of TCX2 and its predicted first neighbors. Given that TCX2 expression has previously been shown to disappear 1-2 hours before stomatal division^12^, we reasoned that we could use our model of TCX2 expression to predict when stem cell division occurs in the root.

**Figure 4.**
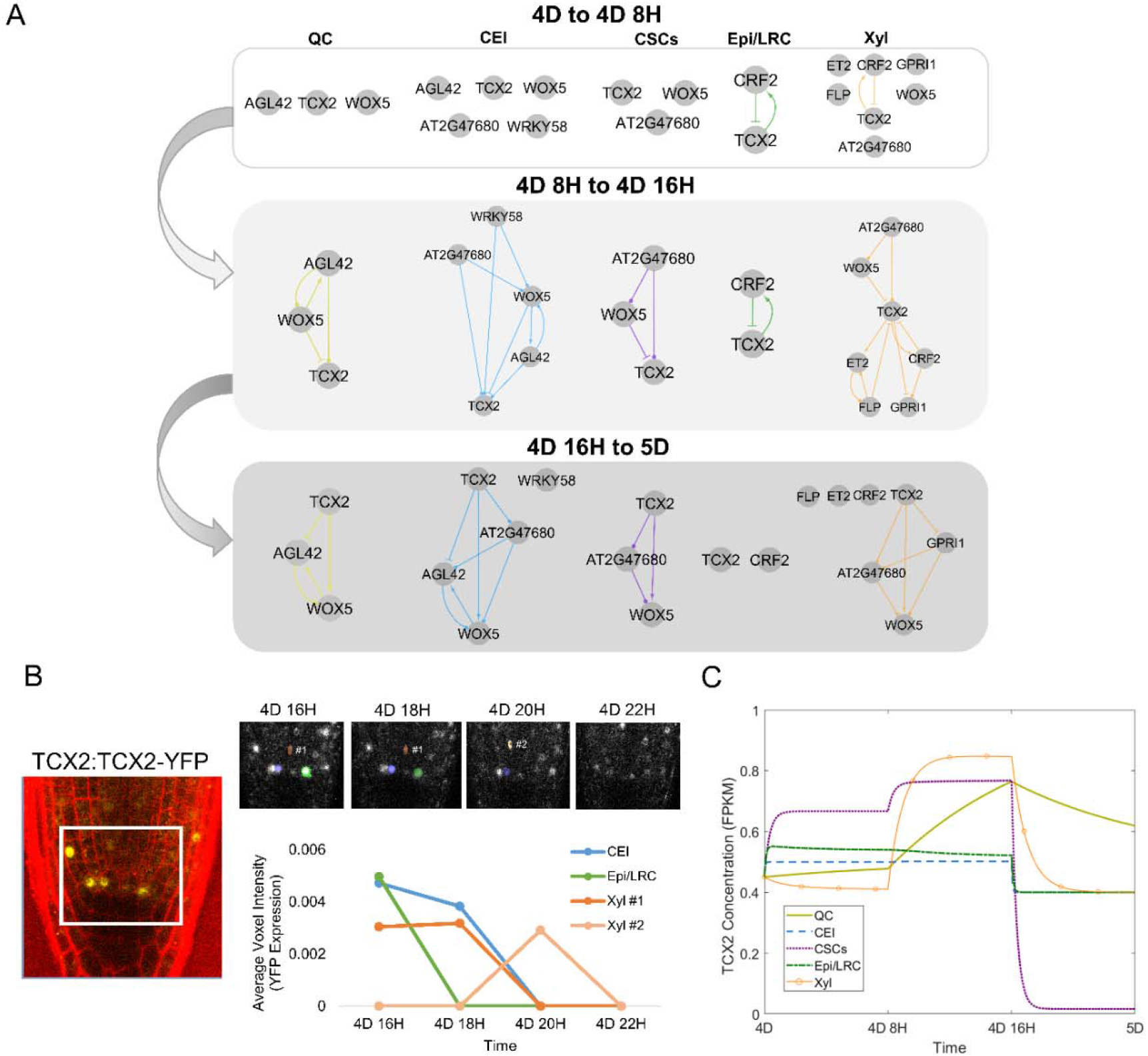
Mathematical modeling of TCX2 network predicts timing of cell division. (A) TCX2 first neighbor TF networks predicted using RTP-STAR on the stem cell time course for 4 day (4D) to 4 days 8 hours (4D 8H) (top), 4D 8H to 4D 16H (middle), and 4D 16H to 5D (bottom). Networks are separated based on the cell type the genes are expressed in: QC (yellow), CEI (blue), CSCs (purple), Epi/LRC (green), Xyl (orange). Arrows represent predicted activation, bars inferred repression, and circles no inferred sign. (B) (left) Representative image of TCX2:TCX2-YFP at 4D 16H. White box represents the stem cell niche were cells were tracked over time. (right, top) YFP-positive cells tracked every 2 hours from 4D 16H (left) to 4D 20H (right). Stem cells that were tracked are marked in blue (CEI), green (Epi/LRC), and orange (Xyl). Two Xyl cells were tracked, #1 and #2. All of these 4 stem cells had no measurable YFP expression at 4D 22H. (right,bottom) Quantification of YFP expression in tracked cells. (C) ODE model prediction of cell-specific TCX2 expression from 4D to 5D. FPKM: fragments per kilobase per million mapped reads.

Our model predicts that there is a significant (fold-change > 1.5) increase in TCX2 expression specifically in the QC and Xyl between 4D 8H and 4D 16H. After this time, our model predicts that the expression of TCX2 in the QC does not significantly decrease and is significantly higher than in all of the actively dividing stem cells (Figure 4C, Supplemental Table 8). Given that the QC is relatively mitotically inactive, this suggests that high levels of TCX2 correlate with a lack of QC division. This prediction is supported by our YFP tracking data which shows that half of the QC cell clusters have either relatively constant or increased TCX2 expression between 4D 16H an 5D (Supplementary Figure 9). Meanwhile, TCX2 expression is predicted to significantly decrease between 4D 16H and 5D in both the Xyl and CSCs, suggesting that these cells divide during this time. This prediction is also supported by our YFP tracking data showing that the majority of Xyl and CSCs cells have low TCX2 expression after 4D 20H (Supplementary Figure 9). In contrast, the CEI and Epi/LRC show only a modest decrease in TCX2 expression between 4D 16H and 5D. This could be due to only some of these cells dividing at that time, as our YFP tracking data shows a large amount of variation in TCX2 expression in these cell populations (Supplementary Figure 9). Taken together, our model and experimental data both suggest that TCX2 not only initiates the division of the actively dividing stem cells, but it also inhibits the division of the QC during the same timeframe, through an unknown mechanism Further, our results allow us to narrow the timing of TCX2-induced stem cell division to a 4-hour window, between 4D 20H and 5D.

## Discussion

Here, we unraveled the communication between stem-cell-specific and stem-cell-ubiquitous networks in the Arabidopsis root through a combination of transcriptomic profiling, GRN inference, biological validation, and mathematical modeling. Our stem cell transcriptional profile revealed that there is both a stem-cell-specific profile that likely provides the foundation for stem cell identity networks as well as a stem-cell-ubiquitous profile that encodes the unique properties shared by all stem cells, such as their ability to asymmetrically divide. Further, our GRN inference predicted that these stem-cell-specific and stem-cell-ubiquitous networks are connected, with the stem-cell-ubiquitous regulators potentially coordinating the downstream stem-cell-specific mechanisms.

Using our network motif score, we identified TCX2 as an important stem-cell-ubiquitous gene that regulates stem cell division by coordinating stem-cell-specific regulatory networks. We validated that TCX2 regulates stem-cell-specific genes through transcriptionally profiling some of the stem cell populations in the *tcx2* mutant, supporting that stem-cell-ubiquitous and stem-cell-specific genes work together to coordinate cell division. Specifically, we were able to validate that 77.78% of the predicted direct targets of TCX2 are differentially expressed in the *tcx2* mutant, 44.54% are predicted in the correct cell type, 58.33% have the correct sign, and 15.08% are directly bound by TCX2. Our results showed that most of the stem cell markers are mis-expressed in other stem cells in the *tcx2* mutant (Figure 3), suggesting that either TCX2 could affect the stem-cell-specific localization of some of these genes or that their cell identity is delayed.

We showed that *tcx2* mutants have additional cell divisions in all stem cell populations, misexpression of known stem-cell-specific marker genes, and higher expression of the cell cycle marker CYCB1;1. Further, our ODE model of the TCX2 GRN illustrated that we can use TCX2 expression to predict the timing of stem cell division. Specifically, our model and TCX2:TCX2-YFP tracking support that a drop in TCX2 expression in most of the stem cell populations between 4D 16H and 5D correlates with stereotypical stem cell division. In contrast, TCX2 levels are relatively stable during this time in the relatively mitotically inactive QC. This is supported by our stem-cell-specific profiling of the *tcx2* mutant which shows that many cell cycle genes, including members of the CYCLIN and CYCLIN DEPENDENT KINASE families, are differentially expressed in different stem cell types (Supplementary Table 5). Notably, TCX2 is a member of the CHC protein family, which in mammalian systems contains components of the DREAM complex such as LIN54^12^. The DREAM complex has been shown to regulate the cell cycle, which supports our proposed role for TCX2 in regulating stem cell division in the Arabidopsis root. It is likely that the other members of the CHC family and other homologs of the DREAM complex in Arabidopsis act together with TCX2 to control this process.

Taken together, our results provide evidence that cell-ubiquitous genes and global signaling mechanisms are important for maintaining stem cell identity and plasticity.

## Materials and Methods

### Lines used in this study

A list of T-DNA insertion lines used in this study is provided in Supplemental Table 9. All T-DNA insertion lines were obtained from the Arabidopsis Biological Resource Center (ABRC: https://abrc.osu.edu/). The marker lines used in this study are described as follows: WOX5:GFP^21^, CYCD6:GFP^22^, J2341:GFP^34^, FEZ:FEZ-GFP^24^, TMO5:3xGFP^25^, CVP2:NLS-VENUS^35^, AGL42:GFP^36^. The TCX2:TCX2-YFP translation fusion is described in^12^, the WOX5:WOX5-GFP translational fusion is described in^33^, the CYCB1;1:CYCB1;1-GFP translational fusion is described in^20^, and the TMO3:TMO3-GFP translational fusion is described in^37^.

### Stem cell transcriptional profile and differential expression analysis

Three to four biological replicates were collected for each marker line. For each biological replicate, 250-500mg of seed were wet sterilized using 50% bleach, 10% Tween and water and stratified at 4°C for 2 days. Seeds were plated on 1x MS, 1% sucrose plates with Nitex mesh and grown under long day conditions (16 hr light/8 hr dark) at 22°C for 5 days. Protoplasting, cell sorting, RNA extraction, and library preparation were performed as described in^36^. For the non-stem cell control, the GFP-negative cells from the AGL42:GFP line were collected. Libraries were sequenced on an Illumina HiSeq 2500 with 100bp single end reads. Reads were mapped and FPKM (fragments per kilobase per million mapped reads) values were obtained using Bowtie, Tuxedo, and Rsubread as described in^38^. Data are available on Gene Expression Omnibus (GEO: https://www.ncbi.nlm.nih.gov/geo/), accession #GSE98204.

Differential expression analysis was performed using PoissonSeq^38,39^. First, stem cell-enriched genes were identified as being enriched (q-value < 0.06 and fold change > 2) in any one stem cell population compared to the non-stem cell control (q-value cutoff of 0.06 was chosen since one of our marker genes, WOX5, had q-value 0.058). Then, genes were classified as enriched in each stem cell type if they had fold change > 2 (enrichment criteria set based on our marker genes) in that stem cell type versus all other stem cell types. If genes were equally expressed in more than one stem cell type, they were considered enriched in multiple stem cell types. All differentially expressed genes are reported in Supplemental Table 1. The Venn diagram in Figure 1C displaying the proportions of genes enriched in each stem cell was constructed using InteractiVenn^40^ (http://www.interactivenn.net/).

### Gene regulatory network inference

The Regression Tree Pipeline for Spatial, Temporal, and Replicate data (RTP-STAR) was used for all network inference. The pipeline consists of three parts: spatial clustering using the *k*-means method^41^, network inference using GENIE3^42^, and edge sign (positive/negative) inference using the first order Markov method^10^. An earlier version of this pipeline was used to infer GRNs of root hair development^43^. This pipeline is implemented in MATLAB and available from https://github.com/nmclark2/RTP-STAR.

For the SCN GRN (Figure 2), networks were inferred for each stem cell separately (resulting in 6 networks, one for each stem cell) and then combined to form the final network. For the stem-cell-specific networks, only the genes enriched in that specific stem cell were used in the network inference. If genes were enriched in multiple stem cells, they were included in all of those individual stem cell networks (e.g. TCX2, which is enriched in all of the stem cells except Protophlo, was included in 5 of the 6 stem cell networks). Genes were first clustered using the mean expression of each gene in each stem cell. Then, network inference was performed using GENIE on only the replicates from that specific stem cell and the SCN marker (e.g. for the QC-enriched cells, only the WOX5:GFP and AGL42:GFP replicates were used). After network inference, the number of edges in the network is trimmed based on the proportion of transcription factors (more transcription factors = more edges kept). Finally, the sign of the edge was determined using a previously published time course dataset of Arabidopsis root stem cells collected from 3 day to 7 day old plants^10^.

For the time point-specific GRNs (Figure 4 and Supplementary Figure 6), we used genes DE in the *tcx2* mutant root tissue sample and enriched in the stem cells. Clustering was performed as for the SCN GRN using mean gene expression in each stem cell. Network inference was performed using the biological replicates from each time point from our stem cell time course collected every 8 hours from 4 days to 6 days old (see Stem cell time course section for more details). Edge sign was determined using this same time course, but using mean expression in all of the time points. One network was built using the biological replicates for each time point and then combined. In Figure 4, the stem cell transcriptomic data was used to determine the stem cell type of each edge.

Due to the pseudo-random nature of *k*-means clustering (i.e., the first clustering step is always random), 100 different clustering configurations (numiter=100 in RTP-STAR parameters) were used for network inference. For the stem cell transcriptional network, edges that appeared in at least 1/3 of the 100 different networks (maxprop=1/3 in RTP-STAR parameters) were retained in the final network as this cutoff resulted in a scale-free network. This parameter was set to edges that appeared in at least 45% of the 100 different networks (maxprop=0.45 in RTP-STAR parameters) for the time point-specific GRNs.

All parameters used to infer these networks in RTP-STAR are included in Supplementary Table 10. All files used to perform GRN inference are available on figshare (see Data Availability section). All network visualization was performed using Cytoscape (http://cytoscape.org/).

### Network Motif Score (NMS)

Five different motifs were used to calculate the NMS namely feed-forward loops, feedback loops, diamond, bi-fan, and multilayer motifs^14–16^ (Supplementary Figure 2). All motifs were significantly enriched in the SCN GRN to a randomly generated network of the same size. First, the number of times a gene appeared in each motif was counted using the NetMatchStar app^44^ in Cytoscape. Then, the counts were normalized to a scale from 0 to 1 and summed to calculate the NMS for each gene. The most functionally important genes are those that have high NMS scores.

### Biological validation

Confocal imaging was performed on a Zeiss LSM 710. Cell walls were counterstained using propidium iodide (PI). Corrected Total Cell Fluorescence (CTCF) was calculated to determine the intensity of cells expressing a fluorescently tagged protein. To complete these measurements, the confocal settings (gain, digital offset, laser percentage) were left constant for the entirety of the experiment. Imaging software (ImageJ) was used to measure the CTCF, which is defined as (Integrated density of GFP)/(Area of selected cells * Mean fluorescence of background) where background is a region of the root with no GFP^45^. The CTCF was divided by the area of the cells (CTCF/area) before performing statistics to account for different numbers of cells selected in each image. When counting cells with GFP expression, a local auto threshold using the Phansalkar method was applied in ImageJ to the GFP channel before counting.

For qPCR, total RNA was isolated from approximately 2mm of 5 day old Col-0, *gata9-1, gata9-2, at1g75710-1, at1g75710-2, rev-5, orc1b-1, orc1b-2, anl2-2* and *anl2-3*, root tips using the RNeasy Micro Kit (Qiagen). qPCR was performed with SYBR green (Invitrogen) using a 7500 Fast Real-Time PCR system (Applied Biosystems) with 40 cycles. Data were analyzed using the ΔΔCt (cycle threshold) method and normalized to the expression of the reference gene UBIQUITIN10 (UBQ10).. qPCR was performed on two technical replicates of two to three independent RNA samples (biological replicates). Differential expression was defined as a p<0.05 using a z-test with a known mean of 1 and standard deviation of 0.17 (based on the Col-0 sample). Primers used for qPCR are provided in Supplementary Table 11. SHR regulation of TCX2 was validated using data from^20^.

### Stem-cell-specific transcriptional profiling in the *tcx2* mutant

Three biological replicates were collected for WOX5:GFP, CYCD6:GFP, FEZ:FEZ-GFP, and TMO5:3xGFP crossed into the *tcx2-2* or *tcx2-3* mutant background. Seedlings were grown and roots were collected as described for the stem cell transcriptional profile. Libraries were sequenced on an Illumina HiSeq 2500 with 100bp single end reads. Reads were mapped and FPKM (fragments per kilobase per million mapped reads) values were obtained using Bowtie, Tuxedo, and Rsubread as described in^38^. Differential expression analysis was performed using PoissonSeq^38,39^. To account for differences in library size between the stem cell transcriptional profile and the TCX2 cell specific transcriptional profile, library sizes were normalized before differential expression was performed. We set a differential expression cutoff of q<0.05 and fold change > 2 based on our cutoff for the stem cell transcriptional profileAllprofile. AllprofileAll differentially expressed genes are reported in Supplemental Table 1.

For the *tcx2-3* transcriptional profile, total RNA was isolated from approximately 2mm of 5 day old Col-0 and *tcx2-3* root tips using the RNeasy Micro Kit. cDNA synthesis and amplification were performed using the NEBNext Ultra II RNA Library Prep Kit for Illumina. Libraries were sequenced on an Illumina HiSeq 2500 with 100 bp single-end reads. Reads were mapped and differential expression was calculated as previously described, except the differential expression criteria were chosen as q<0.5 and fold change > 1.5 based on the values for TCX2, which was assumed to be differentially expressed in its own mutant background.

All differentially expressed genes are reported in Supplemental Table 5. Data for both the stem cell type specific profiling and root tip profiling are available on GEO, accession #GSE123984.

### TCX2:TCX2-YFP and CYCB1;1:CYCB1;1-GFP tracking

Confocal images of the TCX2:TCX2-YFP, CYCB1;1:CYCB1;1-GFP, and CYCB1;1:CYCB1;1-GFP x *tcx2* lines were obtained by imaging roots submerged in agar every 2 hours. A MATLAB-based image analysis software (https://github.com/edbuckne/BioVision_Tracker) was used to detect, segment, and track individual cells expressing YFP/GFP in 3D time-course fluorescence microscopy images^46^. The average voxel intensity, which is a proxy for YFP/GFP expression, was measured as the average voxel value within the set of voxels describing a segmented cell.

### Scanning Fluorescence Correlation Spectroscopy (Scanning FCS)

Image acquisition for Scanning FCS was performed on a Zeiss LSM880 confocal microscope. For Number and Brightness (N&B) on the TCX2:TCX2-YFP and 35S:YFP lines, the parameters were set as follows: image size of 256×256 pixels, pixel dwell time of 8.19 μs, and pixel size of 100 nm. The 35S:YFP line was used to calculate the monomer brightness and cursor size as described in^27,47^. For Pair Correlation Function (pCF) on the 35S:GFP, TCX2:TCX2-YFP and TMO3:TMO3-GFP lines, the parameters were set as follows: image size of 32×1 pixels, pixel dwell time of 8.19 μs, and pixel size between 100-500nm. The movement index (MI) of the 35S:GFP line was used as a positive control. All analysis was performed in the SimFCS software as described in^27,47^.

### Stem cell time course

Two to three biological replicates were collected for each time point. For each biological replicate, 100-250mg of PET111:GFP seed were wet sterilized using 50% bleach, 10% Tween and water and stratified at 4°C for 2 days. Seeds were plated on 1x MS, 1% sucrose plates with Nitex mesh and grown under long day conditions (16 hr light/8 hr dark) at 22°C for 4 days, 4 days 8 hours, 4 days 16 hours, 5 days, 5 days 8 hours, 5 days 16 hours, and 6 days. Roots were collected at the same time of day for all samples to minimize circadian effects. GFP-negative cells were collected as PET111:GFP marks only the differentiated columella, so collecting the surrounding GFP-negative cells results in a population of mostly stem cells. Protoplasting, cell sorting, RNA extraction, and library preparation were performed as described in^36^. Libraries were sequenced on an Illumina HiSeq 2500 with 100bp single end reads. Reads were mapped and FPKM (fragments per kilobase per million mapped reads) values were obtained using Bowtie, Tuxedo, and Rsubread as described in^38^. Data are available on Gene Expression Omnibus (GEO: https://www.ncbi.nlm.nih.gov/geo/), accession #GSE131988.

### Ordinary Differential Equation (ODE) modeling

ODE equations were constructed based on the GRNs shown in Figure 4A. One set of equations was built for each gene in each cell type. The equations changed at 4D 8H and 4D 16H to account for the changes in the predicted network (as shown in Figure 4A). If a sign was not predicted in the network, it was assumed that the regulation was positive (activation) in the model. A schematic showing the location of genes, and what proteins can move between cell types, is presented in Supplementary Figure 7. All equations are provided in Supplemental Equations.

A sensitivity analysis was performed using the total Sobol index^25,38,39^. Sensitive parameters were defined as having a significantly higher (p<0.05) total Sobol index than the control parameter using a Wilcoxon Test with Steel-Dwass for multiple comparisons. (Supplemental Table 6) The sensitive diffusion coefficients and oligomeric states were experimentally measured using scanning FCS. The remainder of the parameters were estimated either directly from the stem cell time course or by using simulated annealing^50^ on the stem cell time course. For simulated annealing, Latin hypercube sampling was used to sample the parameter space for a total of 50 sets of initial parameter estimates. Each set of initial estimates was fit to the residual function using simulated annealing with least squares (simulannealbnd function in MATLAB) for 5 minutes (total runtime = 250 minutes for 50 sets of initial estimates). The average of the 10 parameter values with the lowest error was used in the final model simulation. All parameter values, and how they were estimated, are reported in Supplemental Table 7. All MATLAB files used for the ODE model are available on figshare (see Data Availability section).

### Statistics

For all confocal phenotyping and RICS analyses, a two-tailed Wilcoxon test (for one comparison) or Steel-Dwass with control (for multiple comparisons) was used as some of the data did not follow a normal distribution. All exact p-values, test statistics, and sample sizes are included in Source Data.

## Data Availability

All raw RNA-Seq data and calculated FPKM values are available on GEO, accession #GSE98204, GSE123984, and GSE131988. The Source Data underlying Figure 3 and Supplementary Figures 3, 4, 5, 8, and 9 are provided as a Source Data file. All raw images, the data used for GRN inference, and MATLAB code for the ODE model are deposited on figshare: <u>10.6084/m9.figshare.c.4539071</u>

## Supplementary Information

Supplementary figures, tables, and equations are included in the Supplementary Information PDF.

## Contributions

NMC and RS conceptualized the study and designed the experiments. NMC and APF performed transcriptional profiling. NMC and MAdLB performed differential expression analysis. NMC, ECN, TTN, TBS, and PJS performed biological validation. NMC, ECN, TTN, and PJS collected confocal images. NMC constructed and analyzed the mathematical model. ARS and DCB contributed the TCX2:TCX2-YFP translational fusion. EB and CMW analyzed the YFP tracking data. NMC and RS wrote the paper, and all co-authors edited the paper.

## Acknowledgements

We thank Christian Hardtke and Antia Rodriguez-Villalon for providing the CVP2:NLS-VENUS line. We thank Rüdiger Simon and Barbara Berckmans for providing the WOX5:WOX5-GFP line. We thank Irena Brglez for her assistance with media preparation. We thank Sarah Schuett and the Flow Cytometry and Cell Sorting Laboratory at North Carolina State University (NCSU) for their assistance with cell sorting. Images in this manuscript were generated using the instruments and services at the Cellular and Molecular Imaging Facility (CMIF) at NCSU. Next-generation sequencing was performed by the Genomic Sciences Laboratory (GSL) at NCSU.

## Funding

This work was supported by an NSF GRF (DGE-1252376) awarded to NMC and APF. EB is supported by a GAAN Fellowship in Molecular Biotechnology (grant #P200A160061). ARS was supported by the Donald Kennedy Fellowship and NIH graduate training grant NIH5T32GM007276 to Stanford University. PJS was supported by an Integrated Molecular Plant Systems Research Experience for Undergraduates (IMPS REU) grant awarded to NCSU. DCB is an investigator of the Howard Hughes Medical Institute. Research in the RS lab was funded by an NSF CAREER grant (MCB-1453130) and the NC Agricultural & Life Sciences Research Foundation in the College of Agricultural and Life Sciences at NC State University.

## Notes

#### Summary of Updates

We have made the following major changes to the manuscript: 1) An extended introduction (lines 29-68) and discussion section (lines 287-333) 2) Additional data on tcx2 mutants (lines 163-173 and lines 203-214). 3) More detailed methods throughout the Methods section

