## Supplementary Information for "Stem-cell-ubiquitous genes spatiotemporally coordinate division through regulation of stem-cell-specific gene networks"

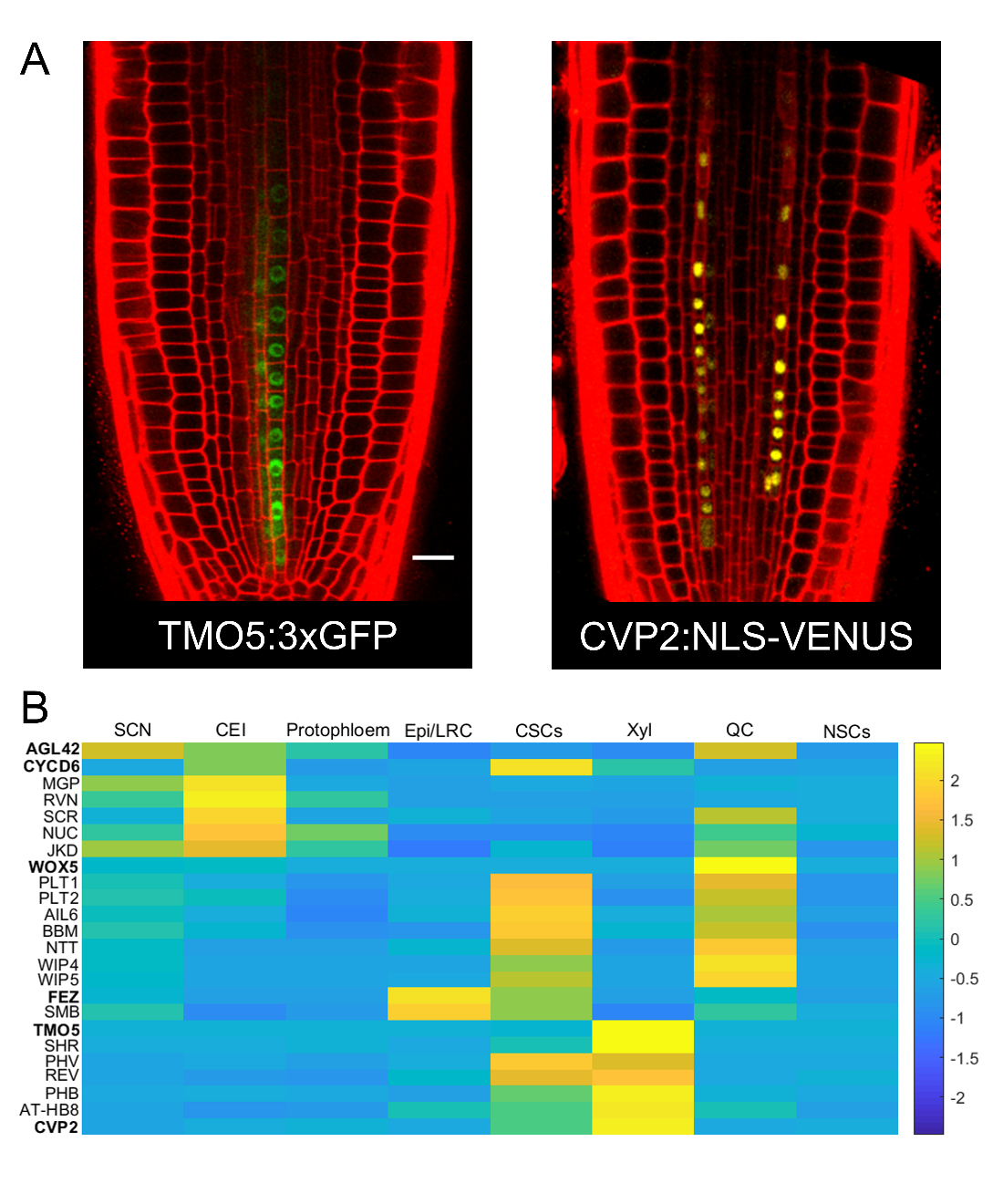
 **Supplementary Figure 1. Expression of known genes in stem cells.** (A) Expression of TMO5:3xGFP (left) and CVP2:NLS-VENUS (right) markers in the meristem. Scale bar = 20µm. (B) Heatmap showing expression of known genes in the stem cells. SCN – stem cell niche; CEI – cortex endodermis initials;; Epi/LRC – epidermis/lateral root cap initials; CSCs – columella stem cells; Xyl – xylem initials; QC – quiescent center, NSCs – non-stem cells. Expression is normalized to mean 0, variance 1 across rows. Color legend represents normalized expression values. Bolded genes are 5 of the 6 stem cell marker genes (the J2341:GFP line is an enhancer trap line and therefore cannot be measured transcriptionally).

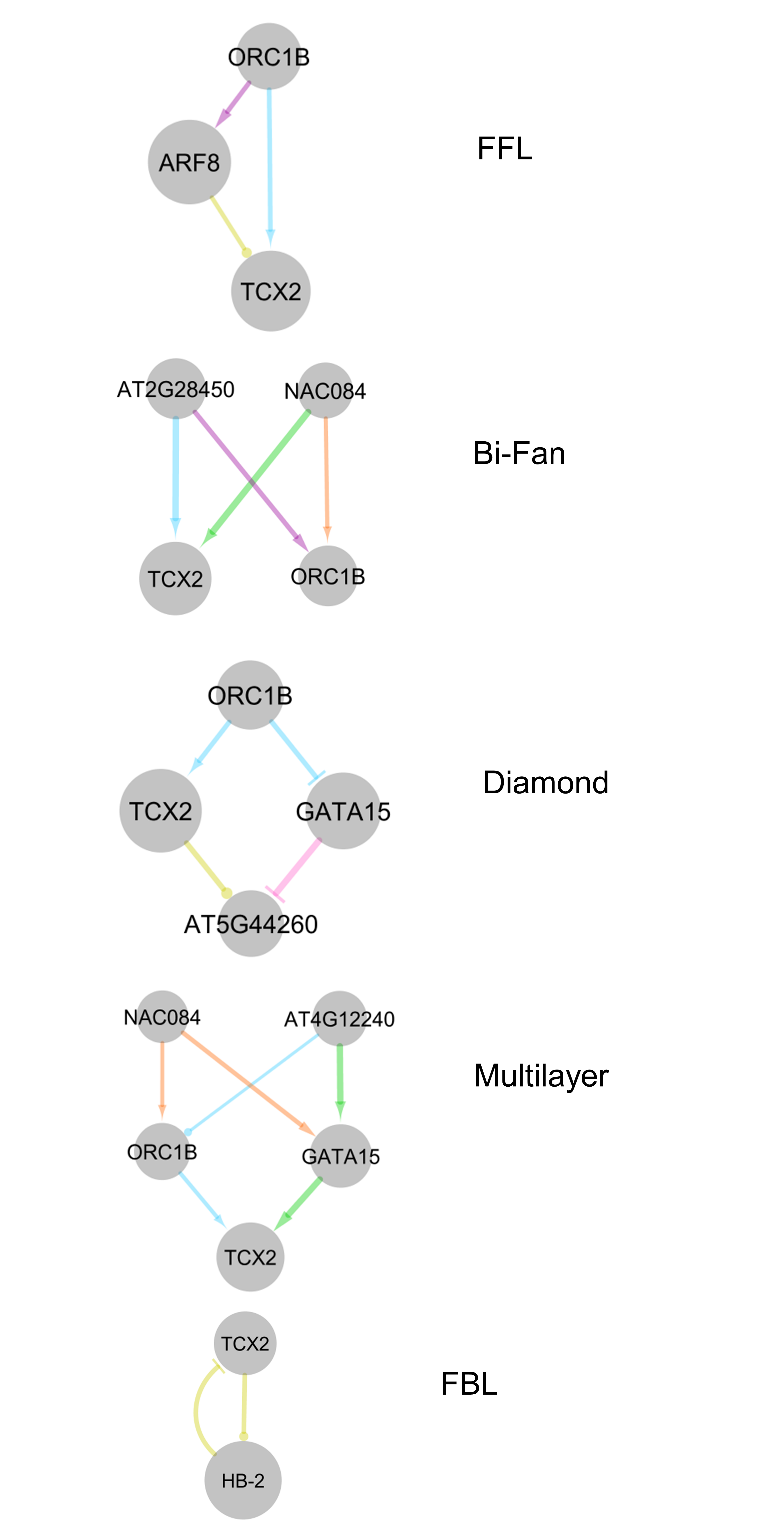

**Supplementary Figure 2. Motifs used to calculate the network motif score.** Examples of the motifs from the stem cell network are shown. FFL – feed forward loop, FBL – feedback loop.

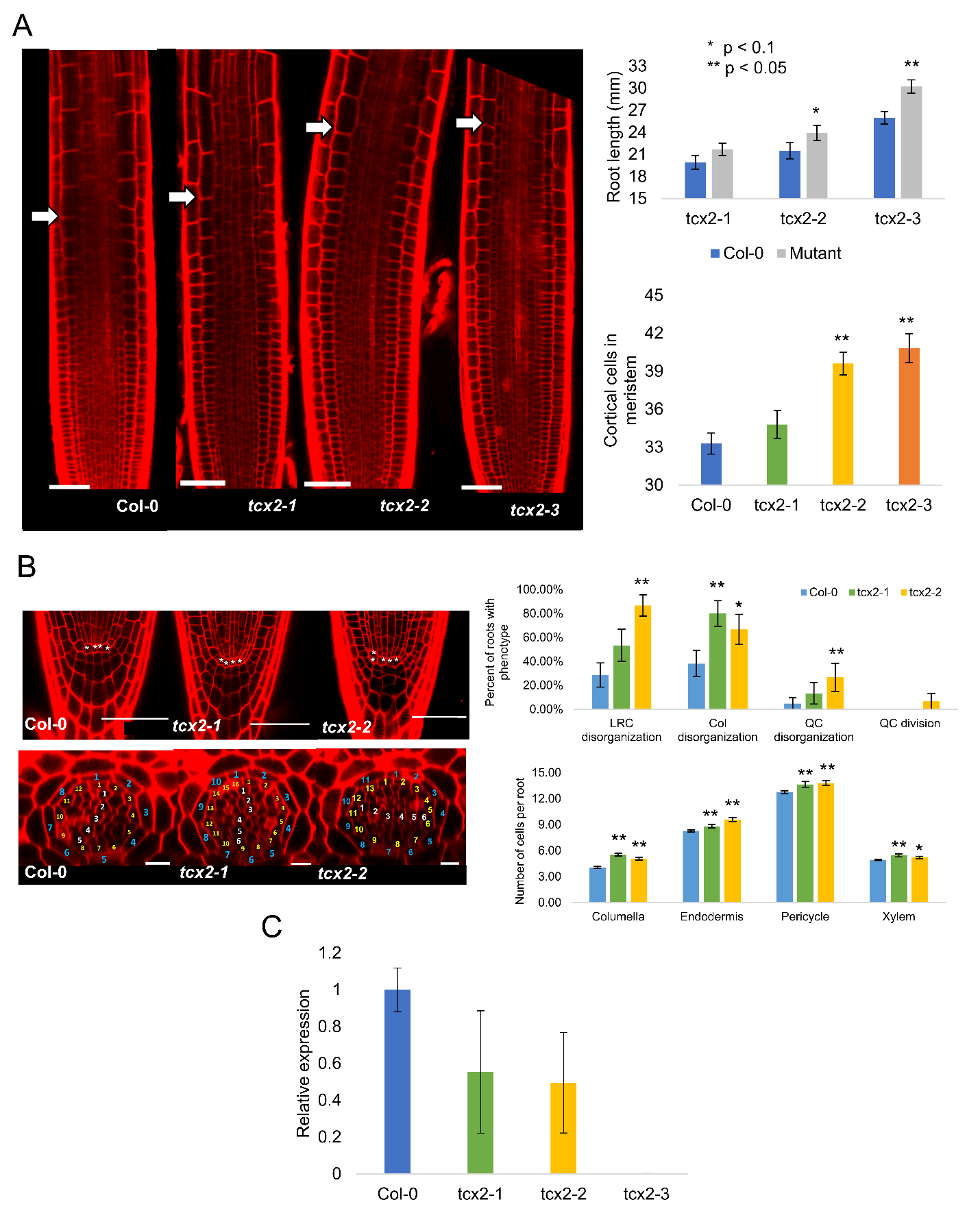

**Supplementary Figure 3. Additional *tcx2* mutant phenotypes.** (A) (left) Representative meristematic zone images of 5 day old WT and *tcx2* mutant alleles. Arrow denotes end of meristematic zone. Scale bar = 50 µm. (right) Quantification of root length after 7 days (top) and number of cortical cells in 5 day old roots (bottom). * denotes p<0.1, ** denotes p<0.05, Wilcoxon Test. Error bars denote SEM. (B) (left) Representative medial longitudinal sections (top) and radial sections (bottom) of WT and *tcx2* mutant alleles. * denotes QC cells. Blue numbers label endodermal cells, yellow pericycle, and white xylem. Top scale bar = 50 µm, bottom scale bar = 10 µm. (right) Quantification of phenotypes. Error bars represent SE. (C) TCX2 expression in WT and the *tcx2* mutant alleles. Averages of 3 replicates are shown. Error bars represent SD. Source data are provided as a Source Data file.

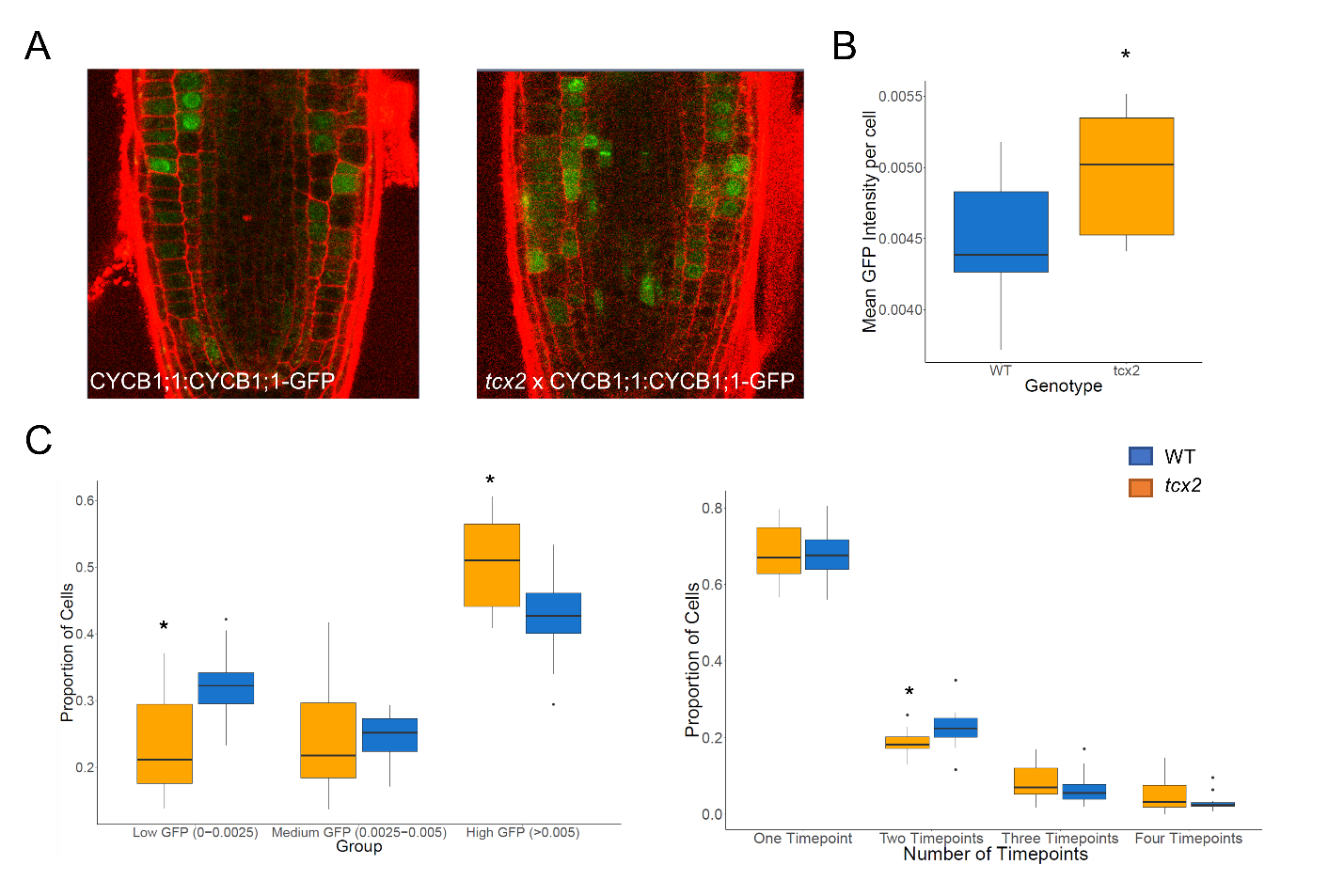

**Supplementary Figure. 4. CYCB1;1 expression in the *tcx2* mutant.** (A) Representative images of CYCB1;1:CYCB1;1-GFP translational fusion in WT (left) and *tcx2* mutant (right) backgrounds. (B) Mean CYCB1;1 expression per cell in WT (blue, n=13 roots) and *tcx2* mutant (orange, n=13 roots). * denotes p<0.05, Wilcoxon test. (C) Proportion of cells with certain GFP levels (left) and that have expression over consecutive time points (right). * denotes p<0.05, Wilcoxon test. Source data are provided as a Source Data file.

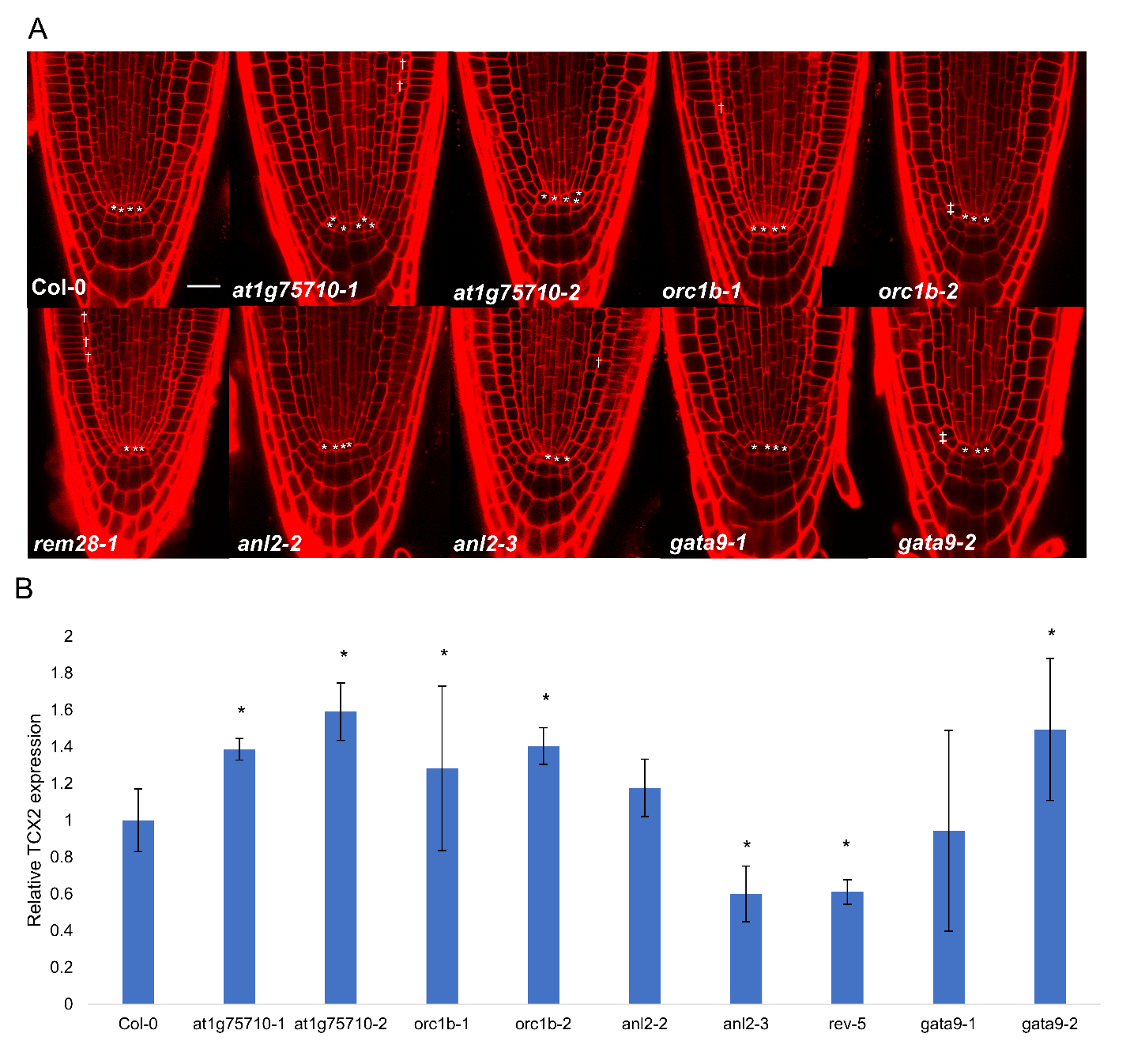

**Supplementary Figure 5. Biological validation of TCX2 regulators and targets.** (A) Medial longitudinal sections of 5 day old Col-0 and mutant plates. * denotes QC, daggers denote extra endodermis divisions, and double daggers denote CEI cells that have not undergone periclinal divisions. Scale bar = 20 µm. (B) qPCR of TCX2 expression in 5 day old Col-0 and mutant roots. * denotes p<0.05, z-test compared to mean and standard deviation of Col-0. Averages of 2-3 biological replicates are shown. Error bars represent SD. Source data are provided as a Source Data file.

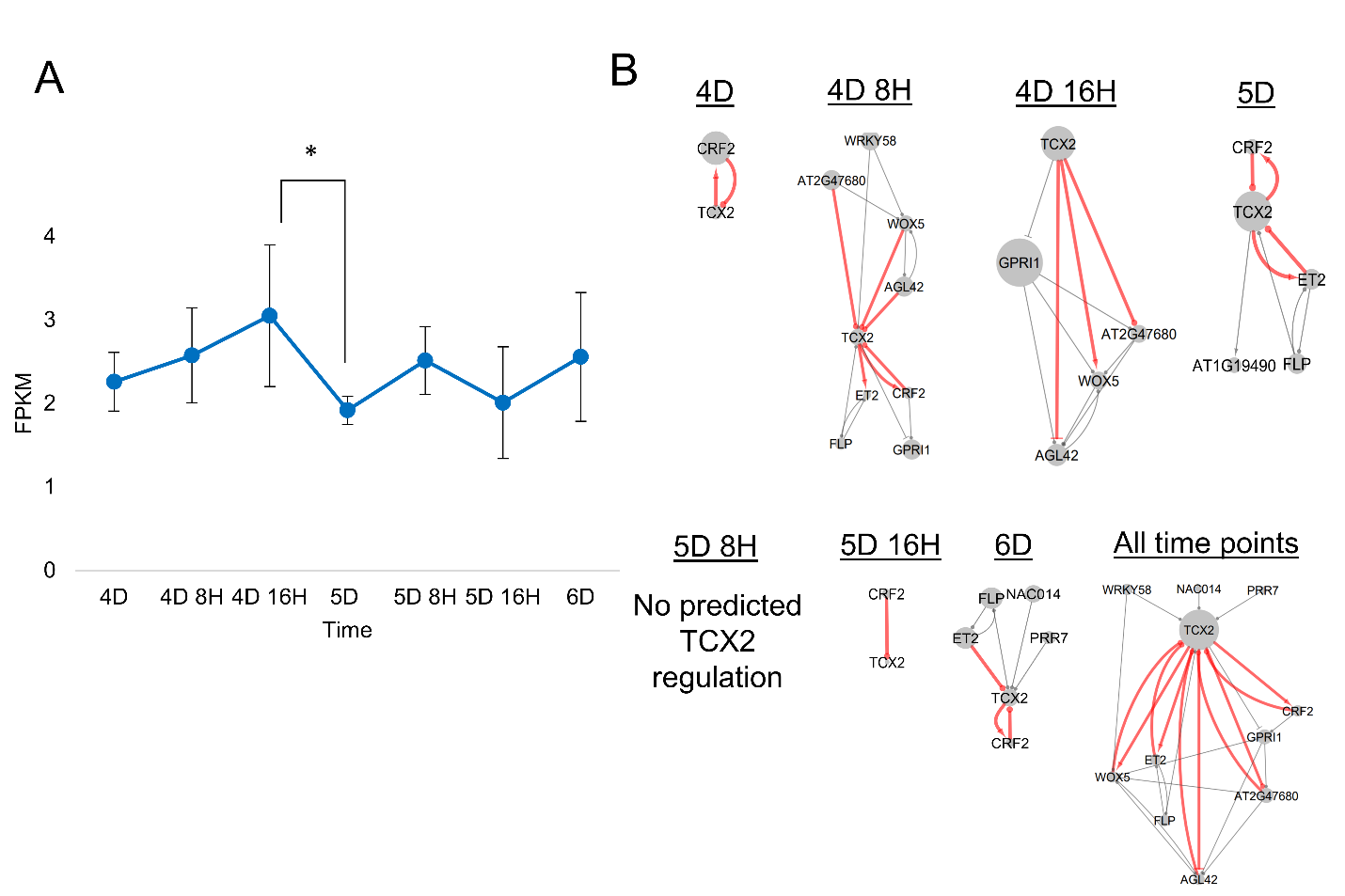

**Supplementary Figure 6. Time-varying TCX2 GRN.** (A) Time course of TCX2 expression in the stem cells. Average of 2-3 biological replicates is shown. * denotes a 1.5-fold change in expression. Error bars are SD. (B) Time-varying first-neighbor TF network of TCX2 in the stem cells. One network is shown for each time point, with the last network (All time points) being the union of the previous 7 networks. Red edges occur at more than one time point, or form feedback loops. Node size represents outdegree in the GRN, not just the TF network shown here. Arrows represent predicted activation, bars predicted repression, and circles no predicted sign.

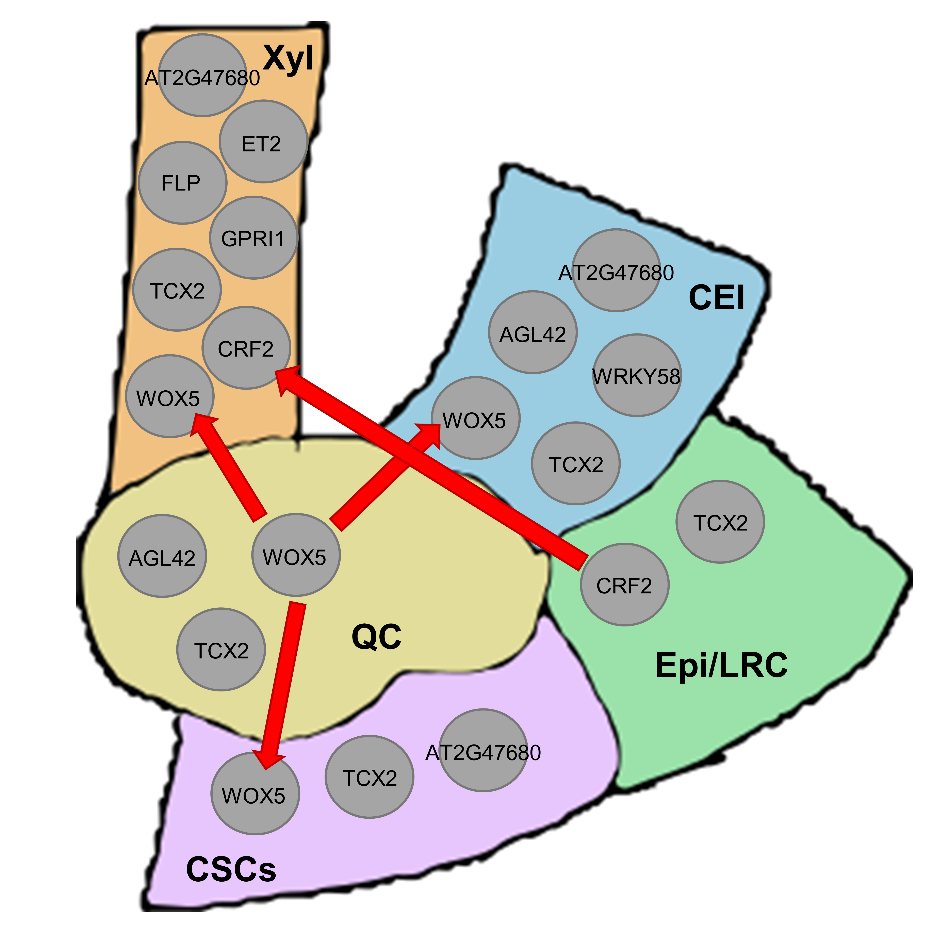

**Supplementary Figure 7. Spatial schematic of TFs involved in the TCX2 ODE model.** Red arrows represent movement of WOX5 and CRF2 protein between cell types.

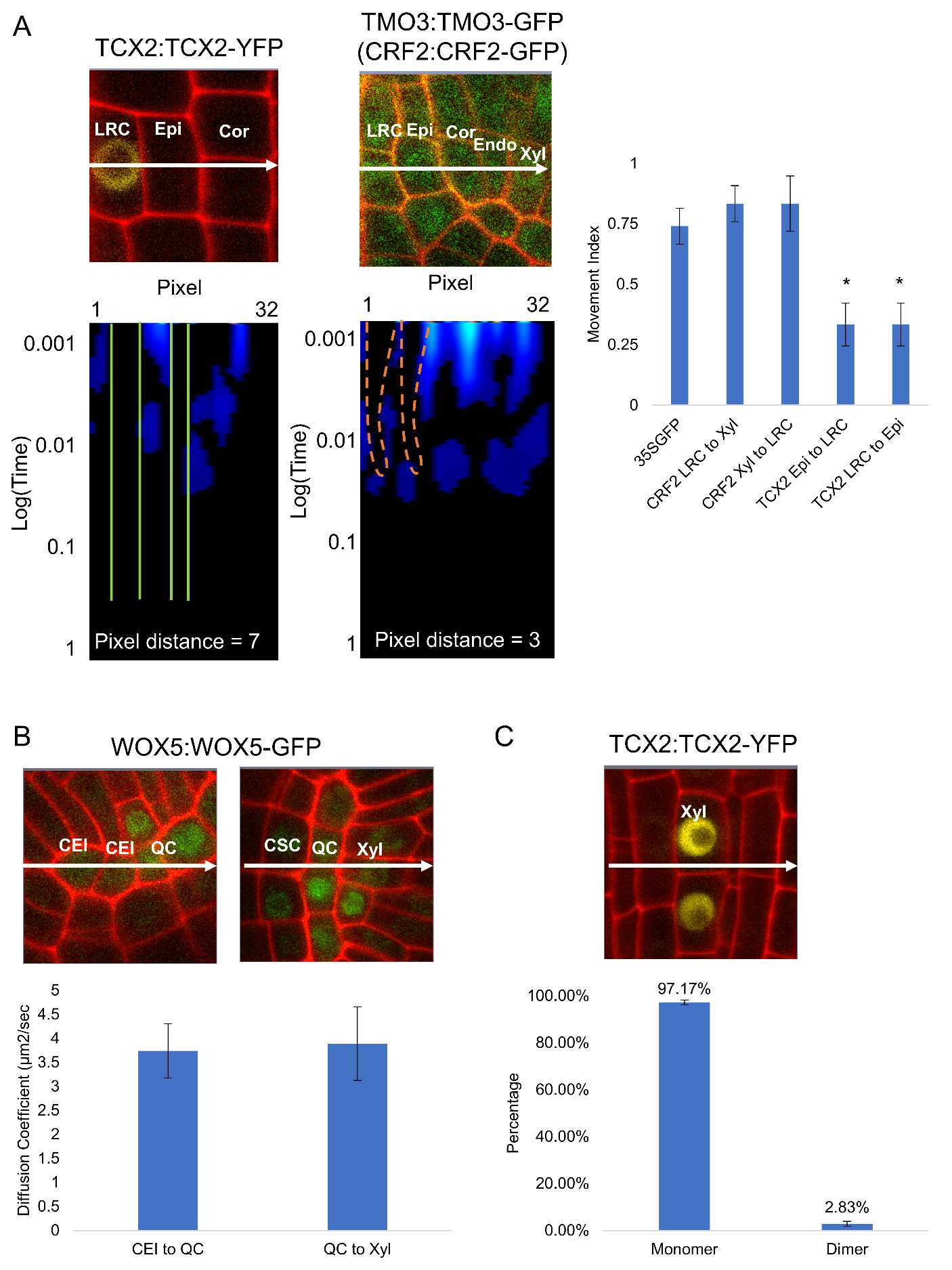

**Supplementary Figure 8. Quantification of protein movement and oligomeric state using scanning FCS.** (A) (left) Representative regions of interest (top) and pCF carpets (bottom) of TCX2:TCX2-YFP and TMO3:TMO3-GFP. Line shows location and direction of the line scan. LRC- lateral root cap; Epi – epidermis; Cor – cortex; Endo – endodermis; Xyl – xylem. Green lines represent no movement, and orange arches represent movement. (right) Quantification of the movement index (MI) (n=6-9). * denotes p<0.05, Wilcoxon test with 35S:GFP as control. (B) (top) Representative regions of interest of WOX5:WOX5-GFP measuring movement from CEI to QC (left) and QC to Xyl (right). CEI – cortex endodermis initials; CSC – columella stem cells; QC – quiescent center. (bottom) Quantification of diffusion coefficient (n=6). Error bars are SEM. (C) (top) Representative region of interest of TCX2:TCX2-YFP measuring oligomeric state in the Xyl. (bottom) Quantification of TCX2 oligomeric state in the Xyl (n=5). Numbers above bars represent means. Error bars are SEM. Source data are provided as a Source Data file.

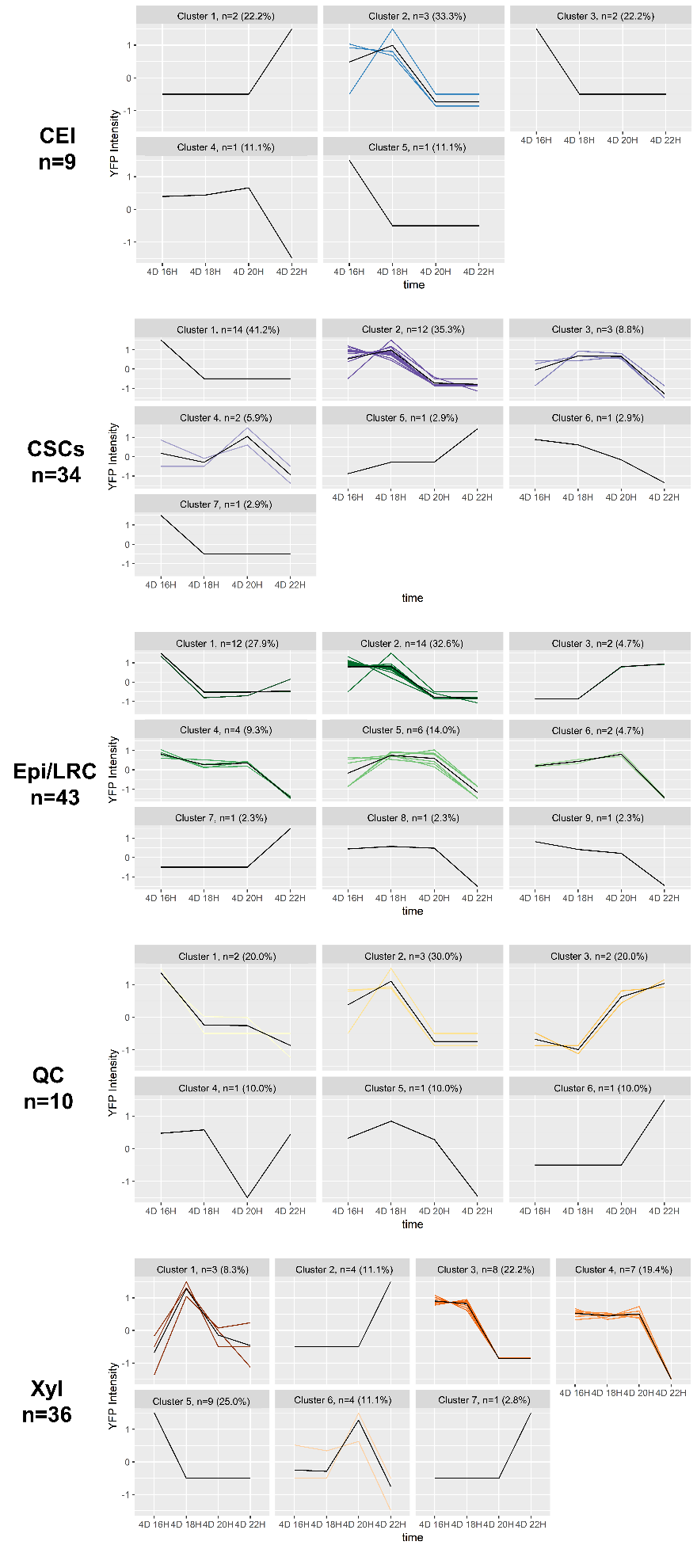

**Supplementary Figure 9. Supervised clustering of tracked TCX2:TCX2-YFP cells.** Supervised clustering of cells tracked across n=21 TCX2:TCX2-YFP roots. Each line represents one cell that was tracked. Numbers in parentheses denote the percentage of tracked cells that are in each cluster. Source data are provided as a Source Data file.

**Supplemental Table Legends
All Supplemental Tables are provided as separate .xlsx files**

**Supplemental Table 1. Differentially enriched genes in stem cells.** Differential enrichment of the 9266 stem cell-enriched genes across the different stem cell types. Enrichment is defined as fold change > 2 and q-value < 0.06 from PoissonSeq versus all other stem cells. Each gene is enriched in the stem cells with a 1 in the corresponding column. The last column is the total number of cells in which each gene is enriched. Cell-ubiquitous genes (red) are defined as genes enriched in 4 or more stem cells, while cell-specific genes (blue) are defined as genes enriched in 3 or less stem cells.

**Supplemental Table 2. Normalized motif scores.** The NMS score and rank are shown for each gene in the network. Only genes with an NMS score > 0 are ranked. Green genes have a known stem cell function and were used to validate the NMS score. The NMS score of TCX2 is highlighted in yellow.

**Supplemental Table 3. Biological validation of TCX2 regulators and targets.** The NMS rank and percentile for each of the 48 genes in the TCX2 first-neighbor TF network are provided. Genes are classified as cell-specific if they are enriched in 3 or less stem cell types. Genes of interest have NMS scores > 0. Mutant obtained: the mutant was obtained and validated in this paper. Phenotype: the obtained mutant did or did not show a stem cell phenotype. Relation to TCX2: the position of the gene in the network.

**Supplemental Table 4. Validation of predicted TCX2 direct targets.** Edges are listed as well as the cell they were predicted in, if they were validated by DAPSeq, if they were validated by RNASeq, if the predicted cell matched at least one of the DE cells, and if the predicted sign matched the known DE sign.

**Supplemental Table 5. Genes differentially expressed in the *tcx2* mutant stem cell types and root tip.** Each tab in the file represents a different pairwise comparison. For the cell type specific data, differential expression was defined as q<0.05 and fold change >2 based on the cutoff for the stem cell transcriptional profile. For the root tip data, differential expression was defined as q<0.5 and fold change >1.5 based on the expression of TCX2 in its own mutant. The 175 genes used in the TCX2 GRN that changes over time are denoted.

**Supplemental Table 6. Sensitivity analysis.** Results for the sensitivity analysis performed on each set of equations (4D to 4D 8H; 4D 8H to 4D 16H; 4D 16H to 5D) are shown. The total Sobol index was averaged across all variables for each of 10 technical replicates. Sensitive parameters are highlighted and have a p-value < 0.05 compared to the denoted control parameter. Yellow parameters were directly estimated from the stem cell time course or experimentally determined using scanning FCS. Green parameters were estimated using simulated annealing on the stem cell time course.

**Supplemental Table 7. Parameter values.** Note that all parameter values are constant across the entire model (4D to 5D). All initial conditions are the average values at 4D from the stem cell time course.

**Supplemental Table 8. Model prediction of TCX2 expression.** FC = fold change.

**Supplemental Table 9. T-DNA lines used in this study.**

**Supplemental Table 10. Parameters used for GRN inference.**

**Supplemental Table 11. qPCR primers used in this study.**
